# Somatic mutation landscape revealed by non-invasive iPSC derivation from urine cells

**DOI:** 10.64898/2026.04.11.717904

**Authors:** Taejeong Bae, Livia Tomasini, Leszek J. Klimczak, Merit Kayastha, Milovan Suvakov, Yeongjun Jang, Alexandre Jourdon, Dmitry A. Gordenin, Flora M. Vaccarino, Alexej Abyzov

**Affiliations:** School of Biosystems and Biomedical Sciences, College of Health Science, Korea University, Seoul 02841, Republic of Korea; Department of Integrated Biomedical and Life Science, Korea University, Seoul 02841, Republic of Korea; Interdisciplinary Program in Precision Public Health, Korea University, Seoul 02841, Republic of Korea; Department of Quantitative Health Sciences, Center for Individualized Medicine, Mayo Clinic, Rochester, MN 55905, USA; Child Study Center, Yale University, New Haven, CT 06520, USA; Integrative Bioinformatics Support Group, National Institute of Environmental Health Sciences, National Institutes of Health, Durham, NC 27709, USA; Mechanisms of Genome Dynamics Group, National Institute of Environmental Health Sciences, National Institutes of Health, Durham, NC 27709, USA; Department of Neuroscience, Yale University, New Haven, CT 06520, USA; Yale Kavli Institute for Neuroscience, New Haven, CT 06520, USA

## Abstract

Somatic mutations that arise post-zygotically create genetic diversity among normal human cells and provide key insights into human development and aging. Fibroblast-derived induced pluripotent stem cells (iPSCs) have proved to be a useful system for disease modelling; however, due to their clonal nature, iPSC lines carry somatic mutations inherited from the founder cells, raising concerns about their genomic integrity. At the same time, this clonality enables single-cell–level discovery of somatic mutations and the reconstruction of developmental lineages. In living individuals, though, this approach requires invasive biopsies and is limited to skin-derived lineages. Here, we generated 33 urine-derived iPSC lines from four males representing two father– son relationships, performed shallow whole-genome sequencing of the lines and analyzed somatic mutations. Derived iPSCs representing single cells from urine carried a few hundred of somatic single-nucleotide variants per genome, dominated by endogenous, clock-like mutational signatures and lacking environmental imprints such as UV-associated mutations. Copy-number analysis identified somatic CNVs in most of the lines and revealed higher CNV burdens in fathers than in sons, consistent with age-related structural mosaicism. Shared mutations across lines enabled reconstruction of cell lineage phylogenetic trees. In summary, urine-derived iPSCs showed genomic alterations comparable to those in fibroblast-derived iPSC lines and represent a valuable non-invasive alternative for disease modeling. Overall, this study provides the first genome-wide characterization of somatic mutations in urine-derived iPSCs and establishes them as a practical and non-invasive platform for charting somatic mutation landscapes and tracing developmental lineages in living humans.

## Introduction

Somatic mosaicism, defined as the accumulation of postzygotic mutations among cells within an individual, is a fundamental feature of human biology ^1,2^. Somatic mutations arise during development and continue to accumulate throughout life, contributing to intercellular genetic diversity (mosaicism) in normal tissues. Over the past decade, analyses of mosaic mutations have been conducted across a variety of human organs, revealing that somatic mutagenesis occurs as a normal consequence of cell proliferation, environmental exposure, and aging ^3–21^. Characterizing such mosaicism provides a window into mutational processes, developmental lineage relationships, and the origins of age-related genomic alterations in healthy individuals ^22,23^.

To achieve single-cell–level resolution of somatic mutations, previous studies have utilized clonal expansion systems, including primary stem cell cultures and induced pluripotent stem cell (iPSC) reprogramming ^5,8,19,21,24^. Among these, skin fibroblast–derived iPSCs have been especially useful, as fibroblasts are easily obtained from living individuals and can be clonally expanded after reprogramming. Our previous works demonstrated that human fibroblast-derived iPSCs harbor about a thousand of somatic single-nucleotide variants (SNVs) at birth, allowing reconstruction of early embryonic lineages and estimation of mutation rates in normal cells ^24–27^. However, skin biopsy to derive fibroblasts is an invasive procedure, even though it is minor and low-risk, and provides access only to the skin, limiting the range of developmental lineage reconstruction.

Besides skin fibroblasts, sources of other accessible somatic tissues, such as peripheral blood, keratinocytes, and urine have also been explored for reprograming ^28^. Cells found in urine derive from the bladder and urinary tract epithelium, as well as from the kidney and ureters, and have a mixed embryonic layer derivation, principally endoderm and intermediate mesoderm ^29–31^. Urine-derived cells offer a fully non-invasive and easily accessible cell source that can be repeatedly collected from living individuals ^32,33^. These cells contain proliferative populations capable of being reprogrammed into iPSCs ^34,35^, making them particularly attractive for studying somatic mutation landscapes in tissues beyond the skin. Similarly to iPSC lines from other tissues, urine-derived iPSCs have been widely utilized to model various diseases, and patient-specific iPSCs have also been generated from urine cells for a range of disorders ^28^. Despite this potential, the genomic characteristics of urine-derived iPSCs and their suitability for reconstructing clonal lineage structures have not been systematically evaluated.

Here, we established iPSC lines from urine-derived cells collected from father–son pairs and performed whole-genome sequencing (WGS) to investigate their somatic mutation landscape. Using an all-to-all (All^2^) variant discovery strategy ^36^ optimized for low-coverage data, we identified somatic single-nucleotide variants and copy-number variations, quantified mutation burdens, inferred mutational signatures, and reconstructed developmental lineages. Our findings demonstrate that urine-derived iPSCs provide a practical and non-invasive system for analyzing somatic mosaicism in living humans, expanding somatic mosaicism studies to cell lineages derived from endoderm and intermediate mesoderm. The somatic mutational burden of urine-derived iPSCs was comparable to that of fibroblast-derived iPSCs, supporting their genomic integrity and highlighting the potential of urine-derived iPSCs for disease modeling and clinical applications.

## Materials and methods

### Collection of urine samples and iPSC line derivation

Informed consent was obtained from each participant in the study according to the regulations of the Institutional Review Board (IRB protocol # 1104008337) and Yale Center for Clinical Investigation at Yale University. Urine samples were collected using the midstream clean catch method. Cells from the urine samples were isolated and expanded following published protocols ^34,35^. The cultured cells were reprogrammed into iPSC lines using a previously published integration-free method ^37^. Briefly, four small-molecule compounds were introduced during the early stages of reprogramming to improve iPSC production efficiency. These include: 1) CHIR99021, a GSK3b inhibitor that promotes reprogramming; 2) A-83-01, a TGF-b/Activin/Nodal receptor inhibitor, which enhances reprogramming when combined with OCT4 and KLF4; 3) Y27632, a ROCK kinase inhibitor that boosts reprogramming efficiency; and 4) PD0325901, a MEK inhibitor that stabilizes the iPSC state. The iPSC lines were propagated using mTESR1 medium (StemCell Technologies) on dishes coated with Matrigel (Corning Matrigel Matrix Basement Membrane Growth Factor Reduced) and propagated using Dispase (StemCell Technologies). In total, 33 iPSC lines were generated from 4 individuals.

### Whole genome sequencing

Genomic DNA was extracted from iPSC lines using QIAamp DNA Mini Kit (Qiagen) following the manufacturer’s instructions. Sequencing libraries were prepared using standard PCR-free methods. Primary urine cells and 24 iPSC lines underwent shallow whole genome sequencing (WGS) at 1X coverage at the Yale Keck DNA Sequencing Facility as an initial genomic QC step. No chromosomal-level aneuploidies were detected in any of the evaluated primary urine cells or iPSC lines. An additional 9 iPSC lines bypassed the 1X WGS QC and were directly submitted to 5X WGS, which likewise did not reveal chromosomal-level aneuploidies. WGS was conducted at 5X or 30X coverage at BGI using 2×100 bp paired reads on a DNBSEQ-G400 sequencer. The FASTQ files were aligned to the human reference genome (GRCh37d5) using BWA. Mark duplicate was performed using PICARD, and indel realignment was conducted using GATK3, resulting in a BAM file for each individual cell line.

### Calling somatic mutations in iPSC lines

We identified somatic SNVs through exhaustive all-to-all comparisons between iPSC lines for each individual ^36^. For each pairwise comparison, we used Mutect2 ^38^. Since Mutect2’s default parameters are optimized for 30X coverage and typically exclude regions with insufficient reference sample coverage in calling process, we changed the value of NLOD parameter to 1.2 to accommodate our shallow (5X) coverage data. This modification allowed inclusion of genomic regions with fewer reference sample reads.

We selected bona fide somatic SNV candidates applying All2 tool ^36^ to the initial calls identified from the exhaustive all-to-all comparisons. The All2 filter was modified to account for the number of cells with no reads at the tested allele, as described below. Candidates were selected if they had a mosaic score above 0.5 and a combined score of 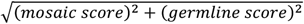 above 0.9 in the All2 filtering step. Additionally, we eliminated calls in inaccessible genomic regions (non-P base region) according to the 1000 Genomes Project mappability mask (i.e., P-bases) and discard germline variants with a population allele frequency of above 0.1% in the gnomAD database.

Using a sample with 30X coverage in individual U11274-03, we compared its somatic SNV call set to the call set of its 5X down-sampled counterpart. The comparison revealed that 5X data tend to call false positive somatic mutations supported by low read counts, which can be mitigated by requiring at least 3 supporting reads for the alternative allele (**Fig. S1**). Based on this analysis, we introduced the *alt3* filter to further improve the accuracy of the calling pipeline for 5X WGS data.

### All2 NRA (No Read Adjustment) mode

The original All2 assumes all cells have data values in any genomic region where it calculates mosaic scores and germline scores, treats all all-to-all pairwise comparisons for any locus as valid, and uses the same fixed cell count, thereby including all cells in the score calculation for all candidate variants (**Fig. S2a**). This assumption does not hold for shallow coverage data, which often contain missing values. Comparisons involving cells with no reads are not valid and introduce biases in the score calculation, leading to the mis-categorization of candidate variants. To address this issue, we modified the All2 algorithm to account for missing values, which vary by locus. The modified version, named NRA (No Read Adjustment) mode, identifies cells with no reads at each locus and excludes them from the score calculation (**Fig. S2b**).

Applying NRA to the 10 cells of individual U11274-03 generated 567 somatic SNV calls, in addition to the 1,854 calls identified by the original All2 algorithm. The additional calls were enriched in loci with a higher number of dropped cells compared to those identified by the original All2 and included 53 miscategorized germline variants, confirmed by their overlap with the father’s germline variants (**Fig. S2c**). Most of default calls identified by the original All2 had sufficient cell counts for accurate score calculation. Additionally, loci with a higher number of dropped cells were more likely to contain miscategorized germline variant calls.

To preserve all the default calls while minimizing false-positive germline variant calls, we only retained NRA-generated calls from loci where more than 5 cells had reads. The excluded call sets under this requirement showed an increased concentration of calls near 50% variant allele frequency in the combined data of all 5X cells, with 11.5% of these calls confirmed as germline by comparison with the father’s germline variants. In contrast, the retained calls contained 5.7% such germline-confirmed variants, reducing false-positive rates by approximately twofold (**Fig. S2d**). We applied NRA to the 13 cells of individual U11274-01 in the same manner.

### Sensitivity estimation and normalized SNV count

To estimate the sensitivity of somatic SNV discovery, we used two approaches. The first approach utilizes germline heterozygous SNPs specific to each individual, identified by comparing the father-son pair, as the reference call set. We identified the reference call sets for the father-son pair in the U11274 family using Mutect2 in paired-sample mode with individual-level pseudo-bulk WGS data, combining all 5X iPSC WGS data.

As in the somatic SNV calling pipeline, we considered only SNPs discovered in confident genomic regions (P-bases in the 1000 Genomes Project genomic mask) and required each SNP call to have at least three alternative allelic reads. Unlike the discovery pipeline, where common SNPs were filtered out, sensitivity estimation relied on rediscovery of germline heterozygous SNPs. For this purpose, we retained common SNPs with a population-wide allele frequency above 0.1% using the gnomAD database rather than filter them out. To eliminate potential homozygous SNPs, we retained only SNPs with a variant allele frequency between 25% and 75% and excluded loci on sex chromosomes. In total, we identified 9,631 and 13,122 individual-specific germline heterozygous SNPs in the father and son, respectively (**Fig. S3a**).

The sensitivity of somatic SNV discovery for each 5X cell was estimated by rediscovering the reference call sets using the somatic SNV calling pipeline by applying all-to-all comparisons of mixed cells, where each cell was compared to other cells from the counterpart (either father or son) within the same family (**Fig. S3b**). The number of the counterpart cells used for mixing was determined to ensure that the total number of cells matched the number of cells within each individual. There are less cells in son than in father. To compensate for the lower number of cells when mixing son’s cells to each father cell, we generated three downsampled 5X data-cells using 30X data from #5 cell.

The other approach utilized 30X data from line #5 and its downsampled 5X data-cells. In this approach, we estimated sensitivity by comparing the call sets from 30X data with those from its 5X downsampled cells. Three trials of rediscovering somatic SNVs (originally identified in 30X data) in 5X data provided not only sensitivity of each downsampled 5X cell but also the numbers of false-positive calls specific to it (**Fig. S3d**). By combining the average count of FP calls estimated by this process with the sensitivities of real 5X data from the other cells derived from the first approach, the normalized SNV count of 5X data was calculated by the formular:

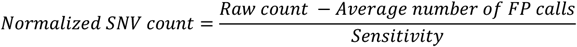

### Mutational signature and mutational motif analysis

For mutational signature and mutational motif analysis, we combined mutations per individual or category. For urine iPSC lines, we aggregated all mutations from 13 and 10 urine iPSC lines derived from two individuals, U11274-01 and U11274-03, respectively. For fibroblast iPSC lines, we used data from a previous study ^24^. We combined all mutations from 25 and 11 fibroblast iPSC lines derived from two individuals, LB and NC0, respectively. For fetal brain and adult brain, we obtained somatic SNVs from previous studies on somatic SNVs in fetal brains ^5^ and adult brains ^39^. We then combined all the SNVs into a single list per category for fetal and adult brains.

To ensure fair comparison, we retained only mutations from previous studies that fell within confident genomic regions (P-base region), as defined by the 1000 Genomes Project mappability mask. This restriction was applied because the mutation-calling pipeline used this study excluded calls in inaccessible genomic regions (non-P base region).

We used an in-house Python script to generate mutational spectra in 96 trinucleotide contexts for urine iPSC lines and fibroblast iPSC lines (**Figure 2a**). Here, we used the combined mutations from all iPSCs within each category. Cosine similarity between the two mutational spectra was computed using the SciPy library. We used SigProfilerExtractor ^40^ (https://github.com/AlexandrovLab/SigProfilerExtractor) to determine the contributions of the known mutational signatures from the COSMIC database in the somatic mutation call sets.

The analysis of mutational motif overrepresentation was conducted using the P-MACD pipeline (https://github.com/NIEHS/P-MACD), which was previously described ^4,20,41–44^. Briefly, the enrichment of defined mutational motifs (**Table S1**) was calculated as the ratio of the observed frequency of a certain mutation within a given mutational motif to its expected frequency, accounting for the surrounding genomic context (±20 bases around the mutated site). The equation used to calculate enrichment is provided below for the mutational motif aTn→aCn as an example, applied to the somatic mutation call sets:

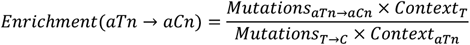

The statistical significance of enrichment values was assessed using Fisher’s Exact test, comparing the mutation frequency within a testing mutational motif to those out of it while accounting for the surrounding genomic context. To correct for multiple hypothesis testing, Benjamini-Hochberg correction was applied to adjust the p-values, ensuring a controlled false discovery rate.

For samples with an enrichment score above 1 and an adjusted p-value ≤ 0.05, the Minimum Estimate of Mutation Load (MutLoadMinEstimate) was calculated to quantify the motif-specific mutagenesis contribution. Using the mutational motif aTn→aCn as an example, this was computed as:

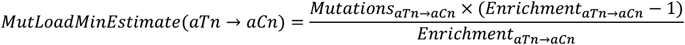

For samples with no statistically significant enrichment, the motif-specific mutation load was set to zero.

### Collecting shared mutations and lineage tree reconstruction

For each individual, a shared mutation matrix was constructed by retaining only mutations observed in multiple cells. For these shared mutations, we genotyped again across all samples starting with the raw call set before applying alt3 filter. By doing so, we included the genotype information for mutation calls with fewer than three alternative allele reads, which had been excluded from the original call sets due to the initial filtering criteria. The final shared mutation matrix contained SNV mutations shared across multiple cells and their corresponding alternative allele counts across all samples.

To identify false-positive somatic calls of germline SNVs in multiple cells, the inclusion of germline variants was tested by analyzing transmitted germline variants, which were identified as germline variants in samples within the same family member. We examined the overlap of shared mutations with the germline SNV call set - using the father’s call set for the son and the son’s call set for the father. The germline SNV call sets were obtained using GATK HaplotypeCaller with default parameters on individual-level pseudo-bulk WGS data, which combined all 5X cells.

Clustering of SNVs and cells was performed using the SciPy library based on binary genotype regardless of the alternative allele read count. Hierarchical clustering was conducted using average linkage with Jaccard distance. A heatmap was generated using the Seaborn library, with different colors representing the count of alternative allele reads.

### Mutation validation using Sanger sequencing

Primers were designed using MacVector v17.0.9 (MacVector Inc) and their specificity was initially determined *in silico* with UCSC Genome Browser (http://genome.ucsc.edu/index.html). Gradient PCR was used to empirically establish the optimal annealing temperature of each primer pair and confirmed primer specify by analyzing PCR products on a 2% agarose gels electrophoresis. PCRs were carried out using 50 ng of genomic DNA per reaction and Phusion High-Fidelity DNA polymerase (Thermo Fisher Scientific) to minimize the polymerase error rate. PCR products were purified using the Mini Elute PCR purification kit (Qiagen). Sanger sequencing was carried out at the Yale Keck DNA Sequencing Facility using standard Applied Biosystems Big Dye chemistry on an Applied Biosystems 3730xL DNA Analyzer. The Sanger sequencing outputs were analyzed in MacVector by visually examining the chromatogram traces.

### Copy number variation detection and analysis

CNVpytor ^45^ was used to detect copy number variations (CNVs) in each cell line through read depth analysis. A non-overlapping bin size of 1000 bp was applied for CNV calling. CNV calls were filtered to retain only those with Q0 values <0.5, indicating that more than half of the reads were uniquely mapped. To exclude low-confidence calls, only those with p-values between 0 and 0.0001 were kept, and calls overlapping regions with >50% unknown bases (i.e., N) in the reference genome were removed (p_N between 0 and 0.5). Only CNV calls ≥10 kb in size were retained for further analysis.

A CNV in a given iPSC line was considered a somatic candidate if its copy number deviated by more than 0.5 compared to the same region in other cell lines. To exclude false positives generated by low-quality mapping in repetitive sequence regions, we calculated the proportion of simple tandem repeats within CNV candidate regions and excluded those with >60% STR bases. Candidate regions were further inspected manually for overlap with segmental duplications and other repeat sequences using the UCSC Genome Browser to ensure high-confidence CNV calls.

Statistical analysis used for CNV sizes and per-cell CNV burdens were performed using the SciPy package in Python. CNV sizes were compared between deletions and duplications using the Mann-Whitney U test. Per-cell CNV burden was compared between fathers and sons using both the Mann-Whitney U test and a two-sample t-test, and Cohen’s d was calculated to estimate effect size. CNV presence versus absence was evaluated using Fisher’s exact test.

### Availability of data and materials

The datasets generated and analyzed during the current study are available in the dbGAP under accession phs004425.v1.p1.

## Results

We collected urine samples from four individuals who share father-son relationships within two families. The collected urine-derived cells were reprogrammed into induced pluripotent stem cells (iPSCs). We established 33 iPSC lines and conducted whole-genome sequencing (WGS) at coverage of 5X, which is required minimum coverage to discover somatic single nucleotide variants (sSNVs) and somatic copy number variations (sCNVs), and at coverage of 30X for a subset of lines (**Figure 1a**). As iPSC lines are typically clonal, we will refer to them below as cells, meaning the founder cells of the lines. By analyzing the WGS data we identified sSNVs and sCNVs in each cell. Profiling these somatic SNVs and CNVs allowed us to estimate mutation burden, reveal mutational signatures, and reconstruct mutation lineages.

**Figure 1.**
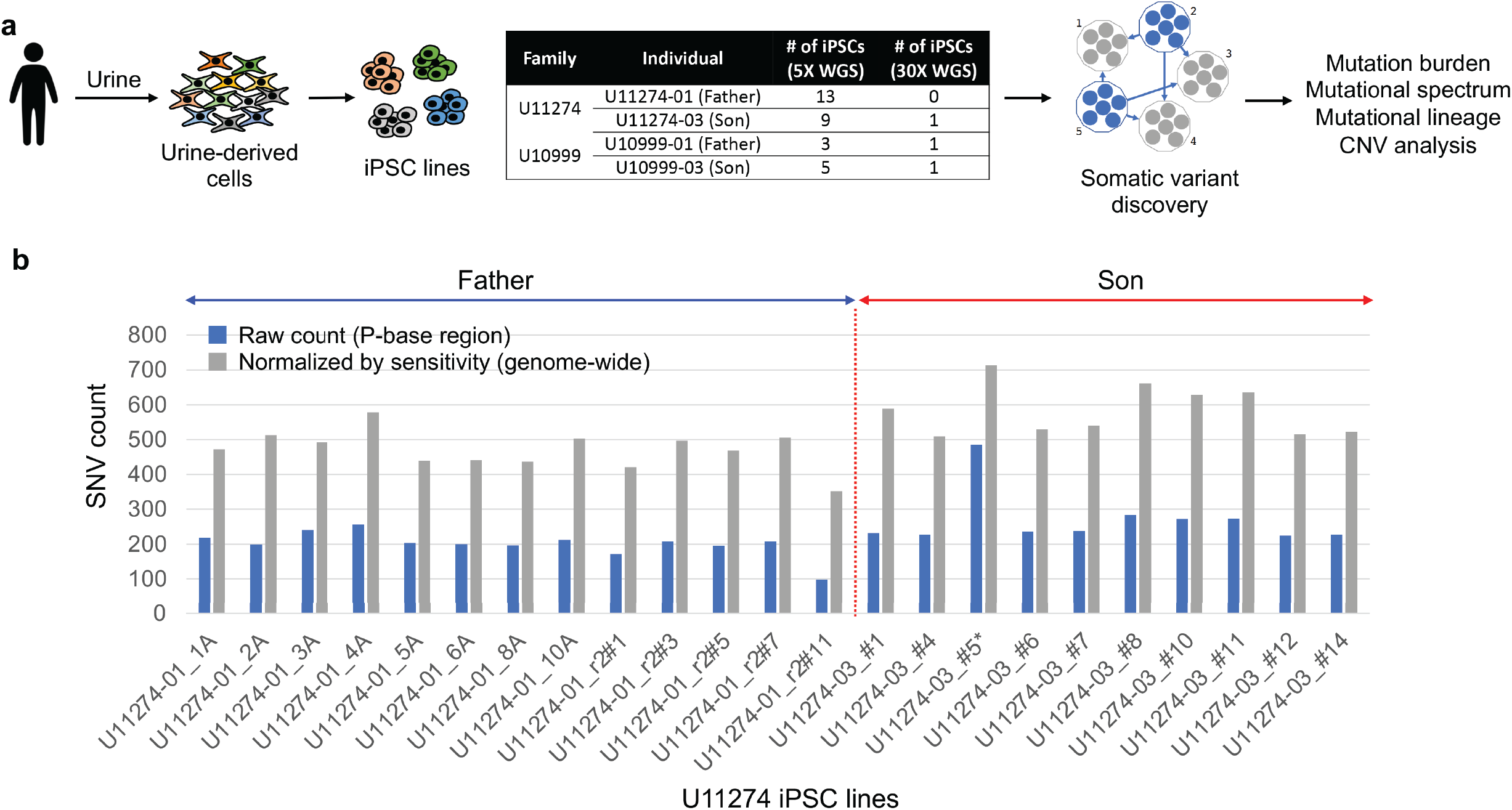
Study overview and mutation burdens. **(a)** Cell cultures were established from human urine samples and cells were reprogrammed into iPSC lines. Whole-genome sequencing was conducted at 5X and 30X coverage. Somatic SNVs and CNVs were identified using the pairwise comparison approach optimized for shallow coverage data, revealing mutational burden, spectrum, and cell lineages. **(b)** Between 98 to 485 SNVs were discovered in 23 urine iPSC lines from the father-son pair in the U11274 family (blue bar). After normalization by the estimated sensitivity and false discovery, mutation burdens were estimated as 352 to 714 per cell (grey bar).

### Mutation burden

Among the two sequenced father-son pairs, only one family (U11274) with a sufficient number of clonally expanded iPSC lines was selected for sSNV discovery and subsequent sSNV analyses. Exhaustive all-to-all (All^2^) pairwise comparison between the genomes of clonally expanded cells is an effective tool to discover somatic mutations in each clone without requiring bulk reference genome data from the same individual ^5,24,25,27,36^. We employed the same comparison approach to discover sSNVs in urine-derived iPSC lines. However, this study presented additional challenges when applying the All^2^ approach due to the shallower sequencing coverage for most iPSC lines, which results in variant allele dropouts, difficulties in distinguishing real mutations from sequencing errors, and contamination of mutation call set by germline variants. To address these challenges, we modified the calling pipeline by relaxing variant filtering during pairwise line comparisons by MuTect2, conducting more stringent filtering of likely germline calls and limiting call set to short read accessible (i.e., P-bases) genomic regions (**Methods, Figure S1** and **S2**). The analysis of a high coverage sample (30X) showed that requiring at least three supporting reads for the alternative allele enables accurate sSNV calling from shallow coverage data (**Figure S1**).

By applying the modified calling pipeline, we discovered from 98 to 485 somatic SNVs per cell line in 23 urine-derived iPSC lines (**Figure 1b**). The sensitivity of sSNV discovery per each iPSC line was estimated based on the recovery rate of germline SNPs unique to each individual by the somatic SNV calling pipeline (**Methods, Figure S3**). The estimated genome-wide sensitivity was 39.7% on average (min 23.7%, max 46.0%, **Table 1**) for the shallow coverage data and 65.9% for the 30X coverage data. Relatively low sensitivity even for high coverage lines was the result of restricting mutations calling to accessible genomic regions, which is ∼71% of the entire human genome (**Methods)**. Normalized by the sensitivity and false discovery rate (**Figure S3, Table 1**), the genome-wide mutation burdens were estimated between 352 to 714 per cell for the 23 urine iPSC lines. The average mutation burden was estimated as 471 sSNVs in the urine iPSC lines from the father and 584 sSNVs in those from the son on average.

**Table 1.** SNV count and sensitivity across all lines.

| iPSC clone | Raw SNV count | Sensitivity<br>observed in Fig. S2c |  | SNV count corrected for FP calls<br>and normalized by sensitivity |  |
| --- | --- | --- | --- | --- | --- |
|  |  | P-base region | genome-wide | P-base region | genome-wide |
| U11274-01_1A | 218 | 61.17% | 43.12% | 332.4 | 471.6 |
| U11274-01_2A | 198 | 50.70% | 35.74% | 361.6 | 513.0 |
| U11274-01_3A | 241 | 65.23% | 45.98% | 347.0 | 492.3 |
| U11274-01_4A | 256 | 59.25% | 41.76% | 407.3 | 577.9 |
| U11274-01_5A | 203 | 60.93% | 42.95% | 309.1 | 438.5 |
| U11274-01_6A | 200 | 59.58% | 42.00% | 311.0 | 441.3 |
| U11274-01_8A | 196 | 58.91% | 41.52% | 307.8 | 436.7 |
| U11274-01_10A | 212 | 55.62% | 39.20% | 354.8 | 503.4 |
| U11274-01_r2#1 | 171 | 52.72% | 37.16% | 296.5 | 420.7 |
| U11274-01_r2#3 | 208 | 55.21% | 38.91% | 350.2 | 496.8 |
| U11274-01_r2#5 | 195 | 54.60% | 38.48% | 330.3 | 468.6 |
| U11274-01_r2#7 | 208 | 54.30% | 38.27% | 356.1 | 505.1 |
| U11274-01_r2#11 | 98 | 33.58% | 23.67% | 248.1 | 352.0 |
| U11274-03_#1 | 231 | 52.16% | 36.76% | 414.8 | 588.4 |
| U11274-03_#4 | 227 | 59.16% | 41.70% | 358.9 | 509.2 |
| U11274-03_#5 | 485 (30X) | 93.52% | 65.92% | 502.9 | 713.5 |
|  | 247 (down-1, 5X) | 49.21% | 34.68% | 472.1 | 669.8 |
|  | 278 (down-2, 5X) | 49.18% | 34.66% | 535.5 | 759.7 |
|  | 268 (down-3, 5X) | 53.02% | 37.37% | 477.8 | 677.8 |
| U11274-03_#6 | 236 | 59.34% | 41.83% | 373.0 | 529.2 |
| U11274-03_#7 | 238 | 58.76% | 41.42% | 380.1 | 539.2 |
| U11274-03_#8 | 283 | 57.63% | 40.62% | 465.6 | 660.5 |
| U11274-03_#10 | 272 | 58.15% | 40.98% | 442.6 | 627.9 |
| U11274-03_#11 | 273 | 57.65% | 40.63% | 448.1 | 635.8 |
| U11274-03_#12 | 224 | 57.66% | 40.64% | 363.0 | 515.1 |
| U11274-03_#14 | 227 | 57.61% | 40.61% | 368.6 | 522.9 |

### Mutational signatures

In the urine-derived iPSC samples, the most frequent mutation type was C to A, whereas C to T was most abundant in skin fibroblast iPSCs from two individuals (LB and NC0) analyzed in a previous study ^24^ (**Figure 2a**). Apart from this difference, we observed similarity between mutation spectra based on 96-trinucleotide motifs of the urine-derived iPSCs and the skin fibroblast iPSCs (cosine similarity of 0.79). To explore the underlying cause of these differences and similarities, the spectrum of combined SNVs were separated into the known mutational signatures of the COSMIC database for each individual. For comparison, we included the mutation spectra of somatic SNVs discovered in fetal and adult brains, which were previously reported as having developmental origin ^5,39^. Eight known mutational signatures were observed across the spectra: SBS1, SBS5, SBS7a, SBS7b, SBS7d, SBS18, and SBS30 (**Figure 2b**).

**Figure 2.**
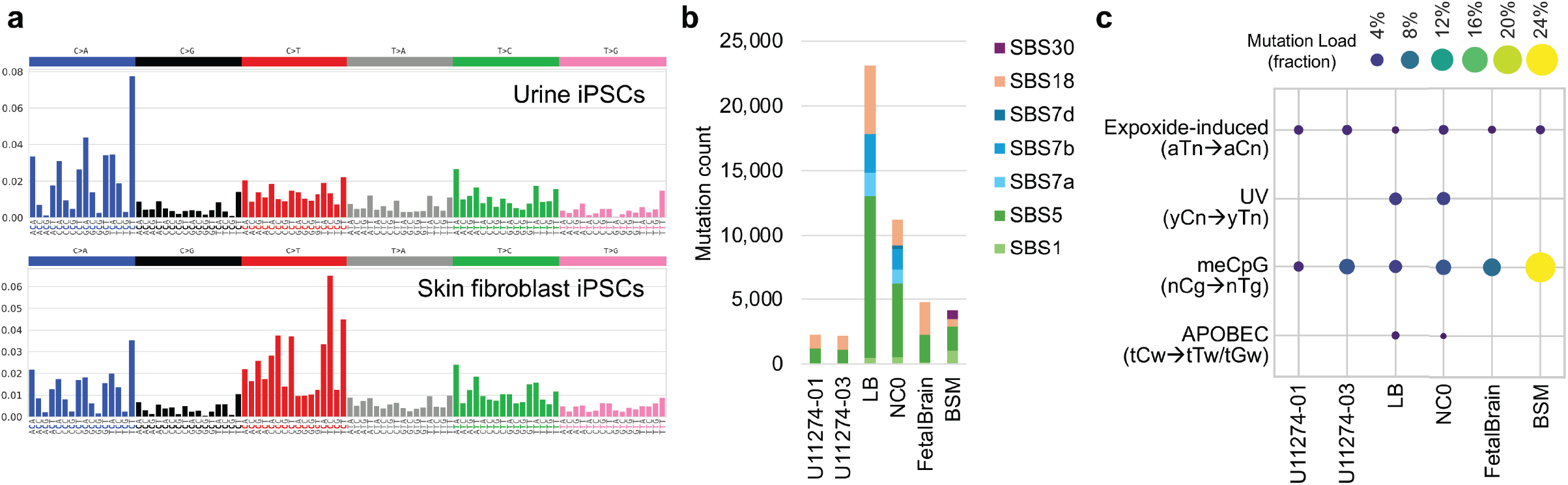
Mutational signatures and motifs. **(a)** Mutational spectra of combined somatic SNVs in urine-derived and the fibroblast-derived iPSCs. The spectra from urine-derived (U11274 combined) and fibroblast-derived (LB/NC0 combined) iPSCs showed a cosine similarity of 0.79. **(b)** Contribution to mutational spectra of the known signatures from the COSMIC database. **(c)** Minimum mutation loads in the mutational motifs by known mutagenesis process. LB and NC0 iPSC lines were derived from skin fibroblasts. U11274-01 and U11274-03 iPSC lines were derived from urine. FetalBrain is the combined somatic SNVs discovered in the single neuronal progenitor cells of the fetal brains (Bae, et al. 2018). BSM (brain somatic mosaicism) is the combined set of somatic SNVs discovered in the adult brains (Bae, et al. 2022). The minimum mutation loads for all 12 mutational motifs are shown in Figure S6, along with their counts and fractions.

The clock-like signature SBS5 was detected at levels between 44% to 54% across all the six mutation spectra (**Figure S4**). In contrast, the clock-like signature SBS1 was observed at low levels, except in adult brains (25%). The enrichment of SBS1 in adult brains is likely attributable to the differences in the discovery methods used. Unlike other call sets, which used clone sequencing capable of detecting single-cell origin mutations, the SNVs in adult brains were identified using high-coverage bulk whole-genome sequencing. Due to its detection limitation, this method can only identify very early somatic mutations that have allele frequencies generally above 1% in tissues and likely enriched for SBS1.

Ultraviolet light exposure signatures (SBS7a, b, and d) were only detected in LB and NC0, the donors of skin fibroblast iPSCs. These signatures were not observed in urine-derived iPSCs and brain samples, consistent with general expectation that only skin fibroblast cells are exposed to UV. The oxidative damage signature SBS18 was observed in all data sets, with higher contributions in cultured cells (iPSCs and clones of fetal brains) as compared to post-mortem bulk brains in the BSM category (**Figure S4**), suggesting that it partially reflects mutations created *in vitro* during cell culture.

SBS30 was detected only in adult brains and is known to be caused by base excision repair deficiency due to NTHL1 inactivating mutations. This observation might be an artifact of signature decomposition rather than representing real mutational processes, possibly attributable to the enrichment of early somatic mutations through the aggregation of SNVs with relatively high allele frequencies across many individuals.

Knowledge-based motif analysis can complement signature-based analysis by highlighting distinct, mechanistically informed mutational motifs. As described previously ^4,20,41–44^, we performed mutational motif-centered analysis for 12 motifs (including 6 sub-motifs reflecting underlying mutational processes) induced by 5 mutational processes (epoxide, UV exposure, methylated CpG deamination, APOBEC enzymatic activity, and redox stress) across different samples (**Table S1**). For four out of five mutational processes (excluding redox stress), the mutational motifs showed statistically significant enrichment after correction for multiple hypotheses testing (**Figure 2c** and **S5**).

Mutation load associated with epoxide-induced mutagenesis (aTn→aCn) was consistently observed at low levels between 1.28% and 2.44% across all samples, suggesting a baseline presence of this mutation type. The methylated CpG deamination motif (nCg→nTg) displayed variable levels across samples, with notable enrichment in adult brains (BSM), indicating active DNA methylation-related processes. UV-associated mutagenesis (yCn→yTn) appeared prominently only in LB and NC0 samples, consistent with their UV-exposed skin fibroblast origin ^4,20^. APOBEC-mediated mutations (tCw→tTw/tGw) were also detected in the LB and NC0 samples at low levels with the mutation load of 1.57% and 0.91%, respectively. It can be explained by marginal APOBEC enzyme mutagenic activities or by unresolvable overlap between yCn UV mutagenesis prominent in skin fibroblasts and the APOBEC motifs tCw (**Figure 2c** and **S6**). Overall, motif-based analysis was largely consistent with signature decomposition analysis and demonstrated the presence of methylation-associated signatures in all samples, as would be expected for endogenous mutagenic processes.

### Lineages of cell ancestries

Somatic mutations shared between cells arise during early development and reveal lineage history ^19,24,25,46^. We identified shared mutations in the two individuals from the same family and provided proof-of-concept lineage reconstruction using iPSC lines (**Figure 3**). To enhance the discovery of shared mutations, we re-genotyped the identified somatic SNVs across all samples and included the genotypes of variants with one or two alternative allelic reads (**Methods**). Low sensitivities due to the shallow sequencing coverage sometime leads to erroneous calling of germline variants as somatic mutations during discovery phase, resulting in enrichment of the shared SNVs across many cells. To address this, we removed ambiguities in the lineage maps by excluding clusters consisting mostly of germline variants (**Methods, Figure S7**). Through this careful procedure, we identified 303 shared SNVs across 13 iPSC lines in the U11274-01 individual and 303 shared SNVs across 10 iPSC lines in the U11274-03 individual (**Figure 3**). Using these shared mutations, we observed six clearly distinguishable lineages in the father and four in the son with a few potential sub-lineages in each individual.

**Figure 3.**
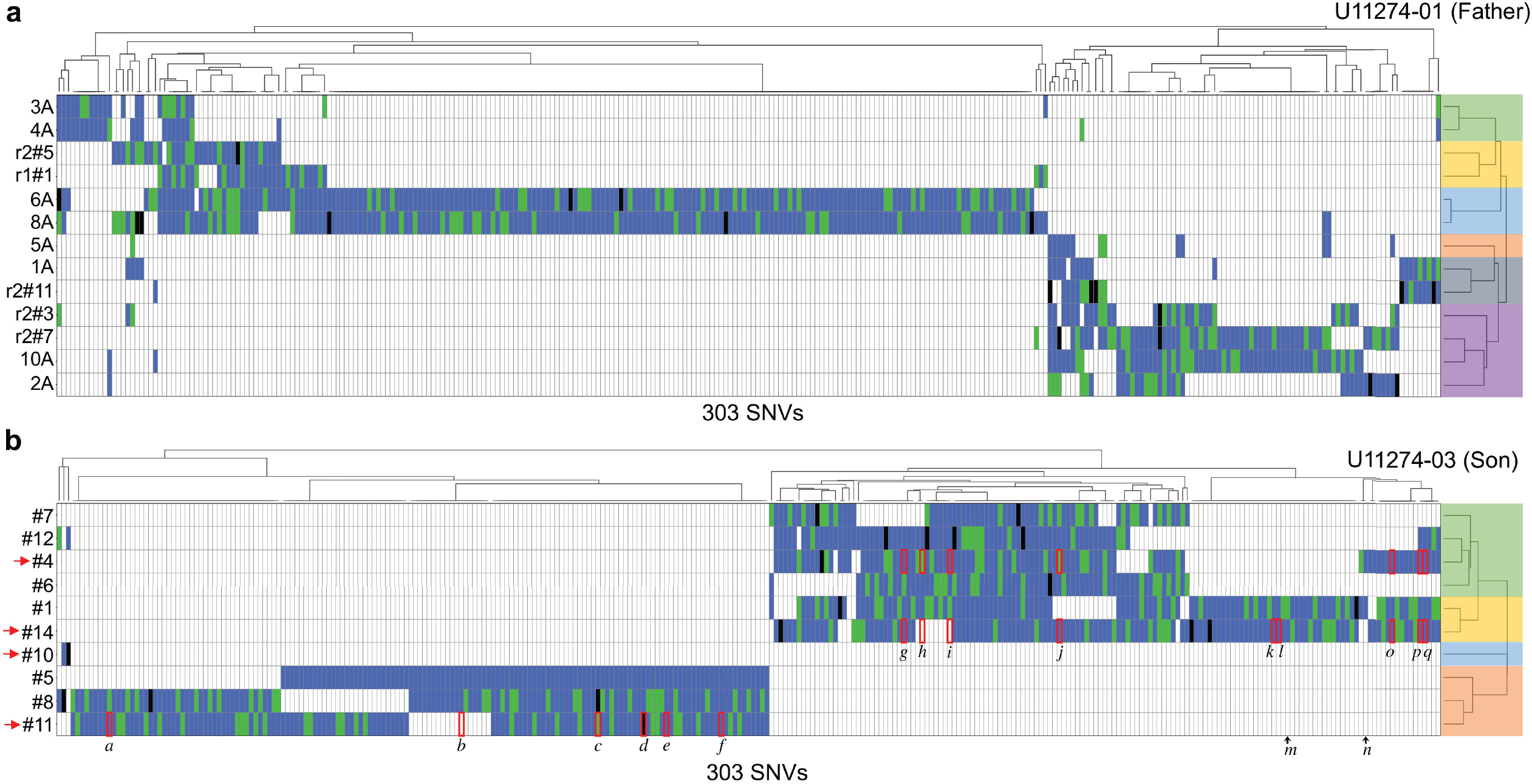
Mutational lineages from iPSC lines grouped by hierarchical clustering of shared SNVs. Two individuals from a family were analyzed: **(a)** U11274-01 (father) and **(b)** U11274-03 (son). Rows represent founder cells of the iPSC lines, and columns represent shared mutations. Blue indicates mutations detected in the discovery phase in cells with at least three supporting alternative allelic reads. Green represents mutations rescued for lineage analysis with two supporting allelic reads, while black denotes mutations rescued with a single supporting allelic read. In B, 15 shared SNVs validated by PCR amplification coupled with Sanger sequencing in the U11274-03 individual are labeled with letters. The two SNVs, m and n, which produced inconclusive validation results, are marked with black arrows at the bottom of the heatmap. The validation experiments were conducted in the four cells (#4, #10, #11, and #14), which are highlighted by the red arrows on the left side of the heatmap. Cells in the heatmap outlined by red denote that the alternative allele was successfully genotyped in the validation experiments. Distinct colors over the lineage trees represent distinguishable lineages among cells—six in the father and four in the son.

To validate the lineage structure in the U11274-03 individual, we conducted genotyping using PCR amplification coupled with Sanger sequencing in four selected iPSC lines representing four lineages. A total of 17 shared SNVs and one additional SNV, which was unique to iPSC line #14, were selected for validation (**Table S2**). Fifteen shared SNVs and a private mutation in cell #14, were successfully validated in the tested lines, confirming that the four lineages were indeed distinct. Notably, three shared mutations—*b, h*, and *i—*were validated in samples that lacked any supporting reads in the original shallow sequencing data, providing evidence consistent with the correctness of the inferred lineage tree. The validation results matched with the expected lineage structure, i.e. mutations were confirmed in lines where they were predicted to be present and absent in lines where they were not.

### Copy number variations

To identify somatic copy number variations, we performed single-cell CNV profiling using CNVpytor and focused on mosaic events present in a subset of cells per individual. Initial candidates identified based on read depth were filtered using multiple criteria, and high-confidence calls were selected after manual inspection (**Methods**). We identified 22 somatic CNVs—including 15 deletions, 6 duplications, and a copy number-neutral loss of heterozygosity (CNN-LOH)—in 16 cells from father-son pairs in two families (**Table S3**). Deletions were the most frequent CNV type, possibly because of higher sensitivity of detecting deletions compared to duplications in WGS-based CNV calling. One deletion and one duplication found in the fathers were shared across two cells (**Figure 4a** and **S8**). Cells 6A and 8A from U11274-01, which shared a deletion, were located within the same lineage branch, as supported by shared somatic SNVs (**Figure 3a**).

**Figure 4.**
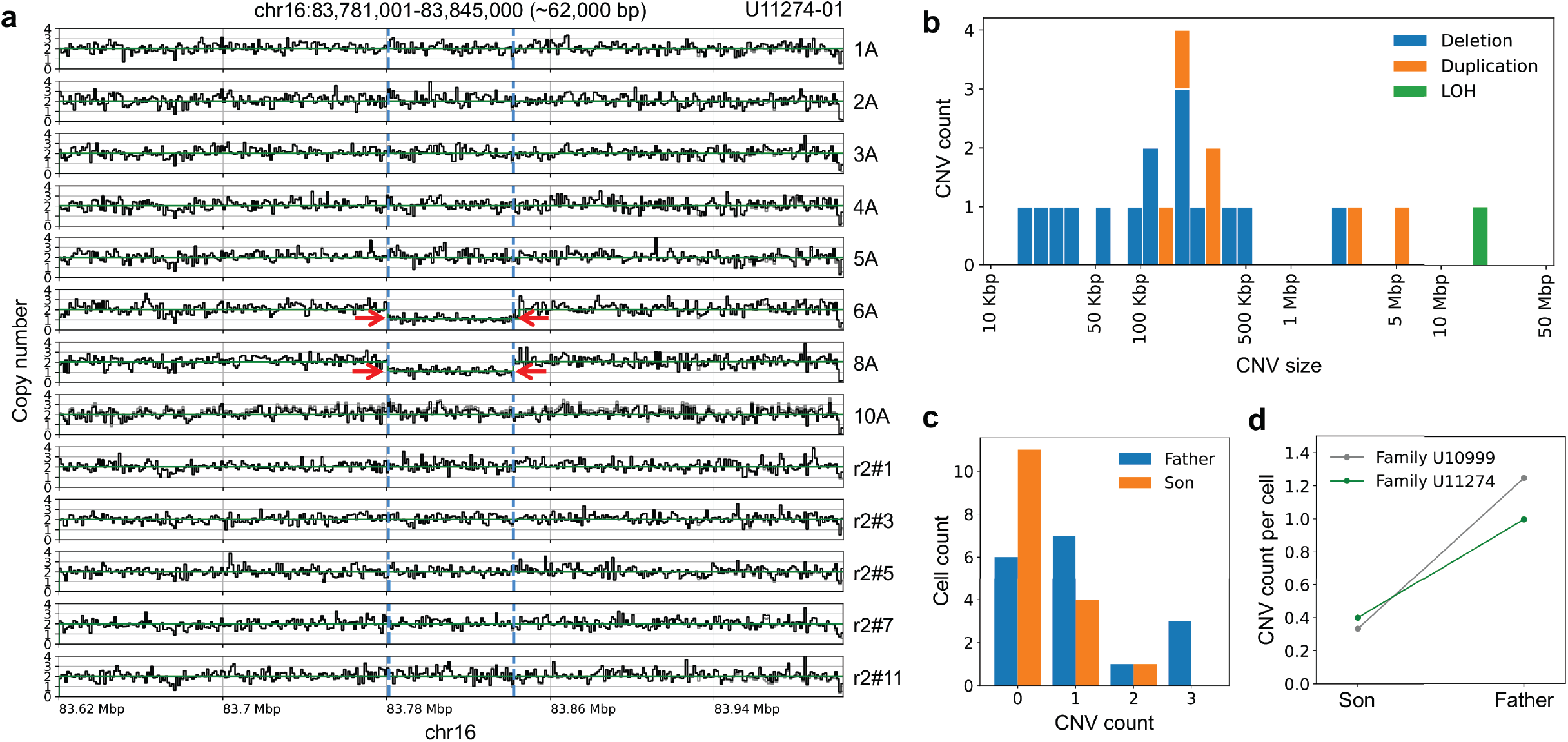
Somatic copy number variations in father-son pairs. **(a**) Shared deletion on chromosome 16 identified in iPSC lines from individual U11274-01 (father of family U11274). Copy number profiles inferred from read depth show a deletion shared by two of the eight cells (red arrows; boundaries marked by blue dashed lines). **(b)** Distribution of somatic CNV sizes across all detected events. Deletions were the most frequent type, with a median size of 123 kb, significantly smaller than duplications (median 309 kb; Mann-Whitney U test, *p* = 0.047). **(c)** Distribution of CNV counts per cell in fathers (blue) and sons (orange) from two families (U10999 and U11274). Fathers showed a significantly higher per-cell CNV burden compared to sons (Mann-Whitney U test, *p* = 0.022; two-sample t-test, *p* = 0.018). **(d)** Average CNV count per cell in each father-son pair. In both families, fathers consistently had a higher somatic CNV burden than sons.

**Figure 5.**
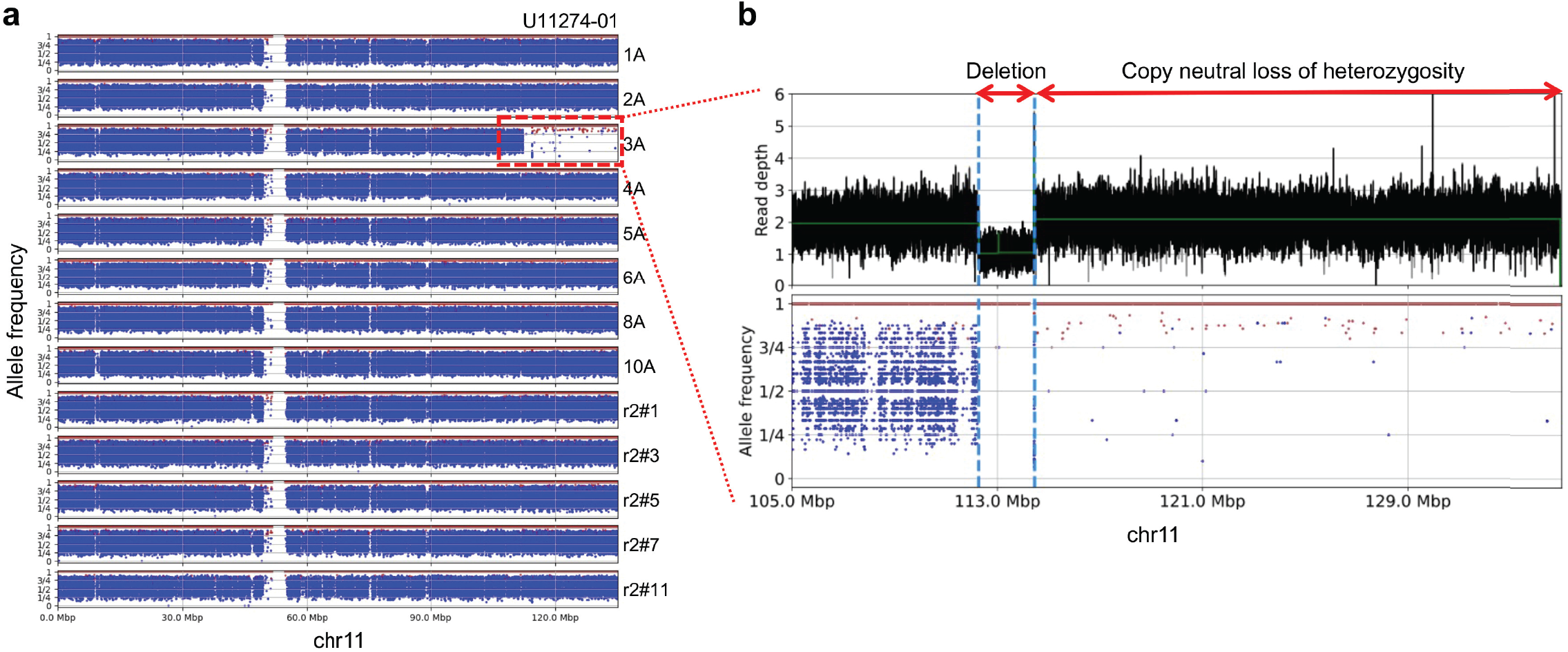
Detection of a deletion followed by a copy-neutral loss of heterozygosity on chromosome 11. **(a)** B-allele frequencies of SNPs across chromosome 11 from multiple single cells (rows) of individual U11274-01. A large loss of heterozygosity region is observed in cell 3A. **(b)** In the highlighted region in cell 3A, a ∼2.2 Mb deletion followed by copy-neutral loss of heterozygosity (CN-LOH) extending to the end of chromosome 11 were identified. The top and bottom panels show copy numbers inferred by read depth and B-allele frequencies of SNPs, respectively.

Event sizes of deletions and duplications ranged from 15 kb to 5.4 Mb (**Figure 4b**). Duplications (median 309 kb) were larger than deletions (median 123 kb; Mann-Whitney test, *p* = 0.047; rank-biserial correlation = 0.58). When randomly pairing deletions and duplications, a duplication was larger than its paired deletion in ∼79% of comparisons. This size difference may partly reflect differences in detection sensitivity between CNV types, as deletions yield stronger read-depth signals, whereas depth gain associated with duplications are more vulnerable to background noise. Distinct mechanisms of CNV generation may also contribute to this size asymmetry. The largest event was a CNN-LOH (∼20.6 Mb) at the end of chromosome 11 in cell 3A from individual U11274-01, which occurred adjacent to a large deletion (∼2.2 Mb) in the same cell (**Figure 4b** and **5**). The adjacent occurrence of the two somatic CNVs within the same cell suggests a single complex rearrangement that may have originated from replication stress or error-prone repair.

Analysis of per-cell CNV burden revealed that the fathers carried, on average, more CNVs per cell (mean = 1.06) compared to the sons (mean = 0.38; **Figure 4c**). Both non-parametric and parametric tests supported this observation (Mann–Whitney U test, *p* = 0.022; two-sample t-test, *p* = 0.018) and a consistent a moderate-to-large statistical difference by Cohen’s d = 0.77. These observations likely reflect longer lifetime of father’s cells leading to accumulation of more CNVs. When considering CNV presence versus absence rather than per-cell CNV counts, fathers again showed a higher proportion of CNV-positive cells (65% vs. 31%). Although this difference did not reach conventional statistical significance (Fisher’s exact test; *p* = 0.057), the trend suggests that somatic CNVs occurred in a larger fraction of father’s cells as compared to son, likely for the same reason – longer lifetime. This was further consistent with family-wise comparisons, in which the average CNV count per cell was higher in fathers than sons in both families (**Figure 4d**).

## Discussion

In this study, we used urine-derived iPSCs from father–son pairs to map somatic SNVs and CNVs, enabling reconstruction of cellular lineages. Mutation burdens per iPSC lines (or cell) ranged from approximately 350 to 710 SNVs per genome, slightly lower but comparable to previous reports from fibroblast-derived iPSC lines, which averaged 850 to 1000 ^4,26,47^. For somatic CNVs, 22 events—including deletions, duplications, and a CNN-LOH—were identified, with CNV burden consistently higher in fathers than sons and the presence of lineage-consistent shared CNVs. These results validate the feasibility of genome-wide mutation profiling from shallow sequencing of urine-derived iPSCs and highlight the importance of read depth-aware filtering in All^2^-based discovery frameworks. Notably, the genome-wide somatic call sets enabled accurate inference of shared mutations across iPSC lines, allowing reconstruction of cellular lineage relationships despite the shallow sequencing depth. The similarity in mutation burdens between urine-derived and fibroblast-derived iPSCs, together with largely comparable mutational spectra, suggests that urine-derived iPSCs maintain genomic integrity at a level similar to fibroblast-derived iPSCs, supporting their suitability for disease modeling and potential clinical applications. This supports the use of shallow WGS of large numbers of clonally expanded lines as a cost-efficient approach for mutation characterization in a sample – an alternative to deep bulk sequencing.

In the analyzed father–son pair, iPSCs from the son had slightly higher SNV counts than from the fathers, which runs counter to the well-recognized monotonic increase in somatic mutation burden with age across human tissues. A possible interpretation is that the father’s iPSC lines may have originated from a restricted developmental lineage—either due to late divergence or clonal expansion—thereby limiting the range of somatic mutations that can be observed in the sampled lines. Because the All^2^ approach depends on contrasting cells’ genomes to each other, such lineage restriction would not find mutations common between all the lines, i.e., those that arose earlier in development or before *in vivo* clonal expansion. Such lineage restriction can lead to an underestimation of the actual mutation burden. Profiling of large somatic CNVs, which rarely happen in development ^48^ and thus represent later somatic events, consistently showed higher CNV frequencies per cell in fathers. Taken together, these observations raise the possibility that the elevated SNV counts observed in the son in the analyzed family reflect sampling of particular lineages during iPSC derivation rather than an apparent inconsistency with age-associated mutational patterns. Future studies incorporating a greater number of independently derived iPSC lineages per individual will be required to more definitively evaluate age-related trends in the SNV burden of urine-derived iPSCs.

Several limitations of this study should be noted. One limitation is that urine-derived iPSCs provide only a partial representation of the somatic lineages present *in vivo*, meaning that the mutation burden inferred from these clones may not fully capture the individual’s overall somatic mutational history. The small cohort size (two families) further limits inference on inter-individual variability and age effects. Another considerable limitation is that shallow WGS of clonal iPSCs yields uneven coverage, reducing overall sensitivity for variant detection and increasing the likelihood of misclassifying germline variants as somatic. Although the optimized All^2^ pipeline mitigates these issues, residual coverage gaps still affect both sensitivity and specificity. Future studies using higher-coverage sequencing and larger cohorts across broader age ranges will improve variant detection and provide a more comprehensive view of the mutational landscape of urine-derived cells.

Together, our study establishes urine-derived iPSCs as a practical and accessible platform for obtaining ancestry and clonal genomic information from living individuals and for interrogating somatic mutagenesis, lineage dynamics, and developmental history in humans. To our knowledge, this represents the first genome-wide characterization of the somatic mutation landscape in urine-derived iPSCs. Looking ahead, extending this framework to include iPSCs derived from multiple tissues within the same individual—such as fibroblasts, blood, and urine—across wider age ranges and genetic backgrounds will deepen our understanding of the origins and tissue specificity of somatic mosaicism throughout the human lifespan.

## Acknowledgements

We thank the family for their participation in this study. We acknowledge Dr P. Ventola and S. Abdullahi for help with subject recruitment. We thank J. Schreiner for the generation of iPSC lines. We thank BGI Americas Corporation for library preparation and whole genome sequencing and the Yale Keck DNA Sequencing Facility for whole genome sequencing and sanger sequencing.

This work was supported by the National Institute of Mental Health (grant no. R01 MH109648 to F.M.V.), National Cancer Institute (grant no. U24 CA220242 to A.A.), National Institute of Diabetes and Digestive and Kidney Diseases (grant no. R25 DK101405 to A.A.), Center for Individualized Medicine at the Mayo Clinic, Korea University (grant nos. K2424621, K2427321, K2427331 and K2503971 to T.B.), the National Research Foundation of Korea (NRF) grant funded by the Korea government (MSIT) (grant no. RS-2025-00515770 to T.B.), US National Institutes of Health Intramural Research Program Project (Z1AES103266 to D.A.G).

## Author Contributions

A.A and F.M.V conceived, designed the study, and edited the manuscript. L.T. and A.J. collected and processed urine samples and established iPSC lines. T.B. and Y.J. processed sequencing data. T.B. generated and analyzed variant call sets, as well as prepared the initial draft of the manuscript and all the figures. L.J.K. and D.A.G. conducted mutational motif analysis. T.B., M.K., and M.S. performed CNV analysis. L.T. conducted validation experiments. All authors read and contributed to the manuscript.

## Supplementary Information

### Supplementary Figures

**Figure S1**. Requiring three supporting reads for mutated bases effectively suppresses false positive calls in shallow sequencing data.

**Figure S2**. Comparison of default and no read adjustment (NRA) modes in All2 filtering.

**Figure S3**. Evaluation of somatic SNV calling sensitivity at 5X coverage using individual-specific germline SNPs from a father-son comparison as a benchmark.

**Figure S4**. Relative contributions of mutational signatures from the COSMIC database.

**Figure S5**. Enrichment of trinucleotide mutational motifs across six different samples.

**Figure S6**. Minimum mutation load estimates of 12 mutational motifs across six different samples.

**Figure S7**. Full version of mutational lineages including ambiguities from iPSC lines grouped by hierarchical clustering of shared SNVs.

**Figure S8**. Shared duplication on chromosome 2 identified in iPSC lines from individual U10999-01.

## Supplementary Tables

**Table S1**. Summary of mutational motifs used in the knowledge-based motif-centered analysis.

**Table S2**. Summary of validation results from experiments using PCR amplification coupled with Sanger sequencing.

**Table S3**. List of copy number variations identified in urine-derived iPSC lines.

