## Supplementary Figures and Tables for "Somatic mutation landscape revealed by non-invasive iPSC derivation from urine cells"

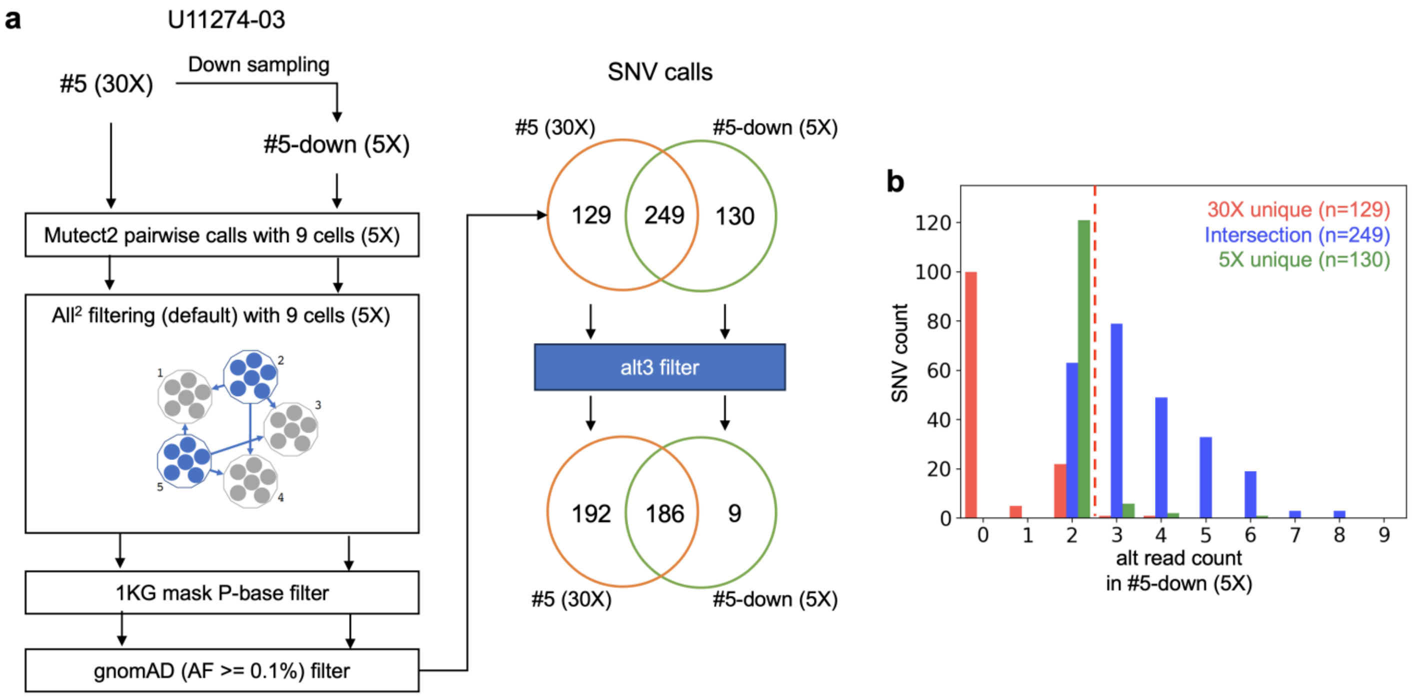


**Figure S1. Requiring three supporting reads for mutated bases effectively suppresses false positive calls in shallow sequencing data. (a)** Schematic overview of the somatic SNV calling and filtering pipeline. We evaluated the pipeline's performance by comparing results from a cell with high-coverage (#5, 30X) and its downsampled counterpart (#5-down, 5X). Pairwise SNV calling was performed using Mutect2 between the test sample and 9 cells (5X), followed by All$^2$ filtering with default parameters. Post-filtering steps included masking inaccessible genomic regions using the 1000 Genomes Project mappability mask (restricting to P-bases only) and excluding from the somatic mutation callset common germline variants with allele frequency above 0.1% in human population. The Venn diagram shows that applying an additional alt3 filter (requiring ≥3 alternative reads) effectively eliminates most of the 5X specific false positive calls. **(b)** Distribution of alternative allele read counts in the 5X dataset for sSNVs uniquely called in 30X (red), shared between 30X and 5X (blue), and uniquely called in 5X (green). The dashed line indicates the alt read count threshold used for alt3 filtering.

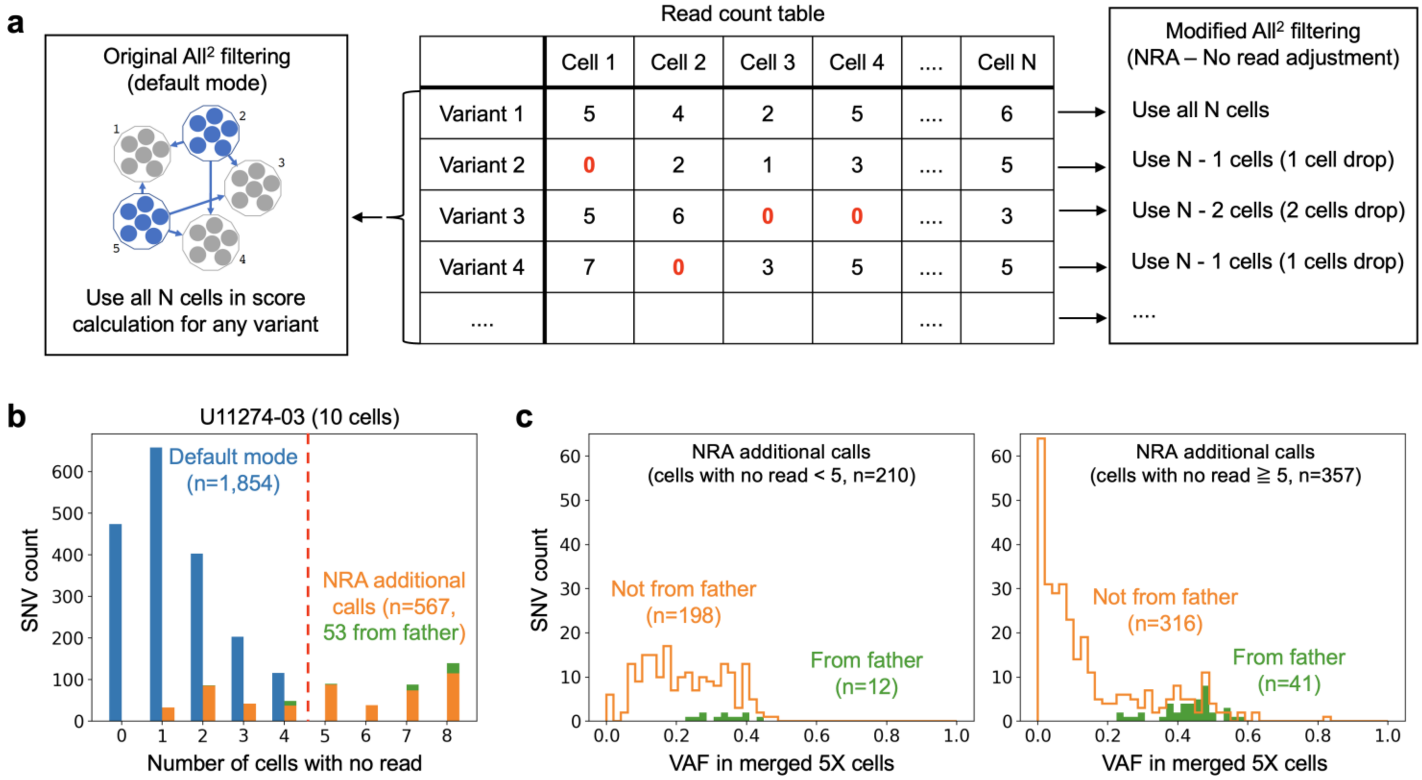


**Figure S2. Comparison of default and no read adjustment (NRA) modes in All$^2$ filtering. (a)** Schematic overview of the differences between default and NRA modes in All$^2$ filtering. In the default mode (left), all *N* cells are included in the score calculation for each variant, regardless of read presence. In the NRA mode (right), cells with no read coverage are selectively excluded from the calculation (e.g., using *N*-1 or *N*-2 cells for different variants), allowing for more flexible handling of missing data. The read count table (center) illustrates how missing read information (red zeros) is treated differently in each mode. **(b)** Distribution of single-nucleotide variant (SNV) counts by the number of cells lacking read coverage in individual U11274-03 (10 cells). The NRA mode identified additional SNVs (orange) by avoiding score calculation distortion from missing data while retaining all SNVs originally called by the default mode (blue). All SNVs identified in the default mode had fewer than 5 cells with no reads. As the number of missing cells increases, a greater proportion of the additional SNVs (green) overlap with paternal SNVs, suggesting contamination with germline SNPs. A threshold of 5 or more cells with no reads (indicated by the red dashed line) was used to filter out less confident additional calls. **(c)** Variant allele frequency (VAF) distributions for NRA-specific calls, stratified by the number of cells without read coverage. Left: variants with fewer than 5 cells lacking reads. Right: variants with 5 or more cells lacking reads. Colors indicate paternal origin of called SNVs (orange: not from father; green: from father).

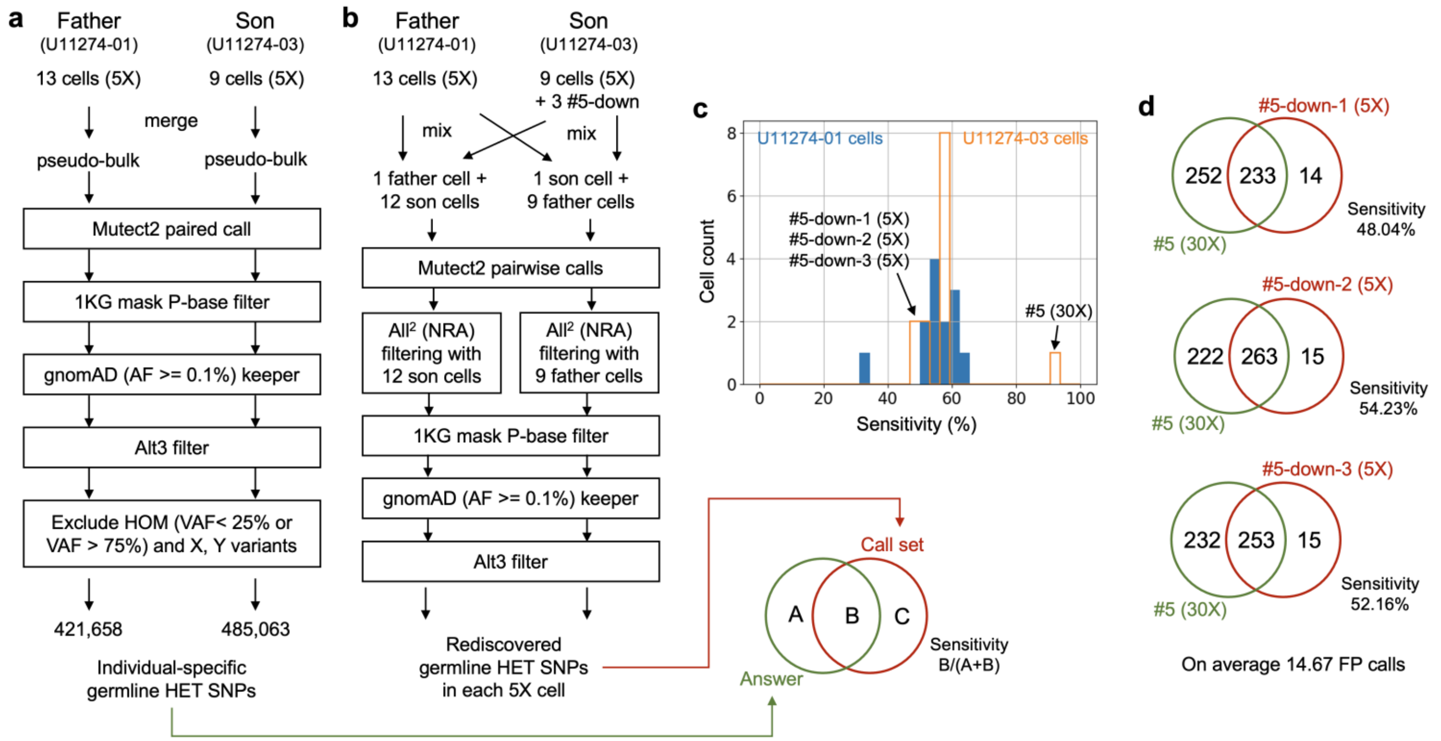


**Figure S3. Evaluation of somatic SNV calling sensitivity at 5X coverage using individual-specific germline SNPs from a father-son comparison as a benchmark. (a)** Identification of individual-specific heterozygous (HET) germline SNPs from father (U11274-01) and son (U11274-03) samples. Pseudo-bulk samples were generated by merging 5X coverage data from 13 (father) or 9 (son) single cells. Mutect2 paired calling was performed, followed by several post-filtering steps: application of the 1000 Genomes Project mappability mask (requiring P-bases), retention of gnomAD common SNPs (allele frequency ≥ 0.1%), and application of the alt3 filter (requiring ≥3 alternative reads) as well as exclusion of homozygous variants (VAF < 25% or > 75%) and variants on sex chromosomes. As the sensitivity calculation was based on individual-specific germline HET SNPs, we included only gnomAD common SNPs in order to retain high quality germline HET calls. The resulting individual-specific HET germline SNPs were used as answer sets for benchmarking. **(b)** Rediscovery of individual-specific HET germline SNPs in each cell from mixing experiments using father and son samples. To assess the sensitivity of identifying true germline HET SNPs from low-coverage data by the somatic SNV calling pipeline followed by All² filtering, we performed pairwise SNV calling with Mutect2 between each cell from one individual and cells from the other individual. The number of selected cells from the opposite individual was chosen to match the total number of cells in the individual being analyzed. The father had thirteen 5X cells, while the son had nine 5X cells and one 30X cell. To match the father's cell count, three additional 5X cells were generated by down-sampling the son's 30X cell. All filters in the pipeline were applied consistently as during discovery (**Fig. S1**), except for the retention of gnomAD variants with allele frequency ≥ 0.1%, which was intentionally included in this analysis. **C)** Distribution of sensitivity within the P-base regions of the 1000 Genomes Project accessible mask. Sensitivity for each cell was calculated as the proportion of rediscovered SNPs (Panel B) among the individual-specific germline HET SNPs (Panel A). The blue-filled histogram represents cells from the father (U11274-01), and the orange-outlined histogram represents cells from the son (U11274-03). **(c)** Sensitivity based on the rediscovery of somatic SNV calls identified in the 30X cell using three down-sampled 5X cells. Alternatively, we estimated the sensitivity of somatic SNV calling in low-coverage data using 3 down-sampled 5X datasets generated from the son's single 30X cell. The sensitivities of these three cells were comparable to those observed in the panel C. This approach also allowed us to estimate the number of false-positive calls, defined as additional SNVs identified only in the 5X cells. The average number of false-positive calls was 14.67.

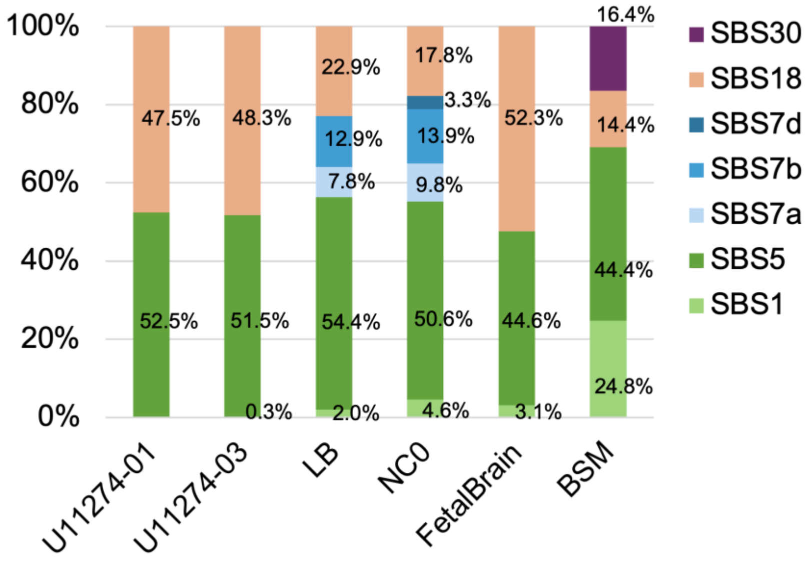


**Figure S4. Relative contributions of mutational signatures from the COSMIC database.** U11274-01 and U11274-03 iPSC lines were derived from urine. LB and NC0 iPSC lines were derived from skin fibroblasts. FetalBrain represents combined somatic SNVs identified in single neuronal progenitor cells from fetal brains (Bae, et al. 2018). BSM (brain somatic mutations) represents combined somatic SNVs identified in adult brains (Bae, et al. 2022). The distinct mutational signature profiles reflect differences in detection method, developmental origin, tissue type, and mutational processes.

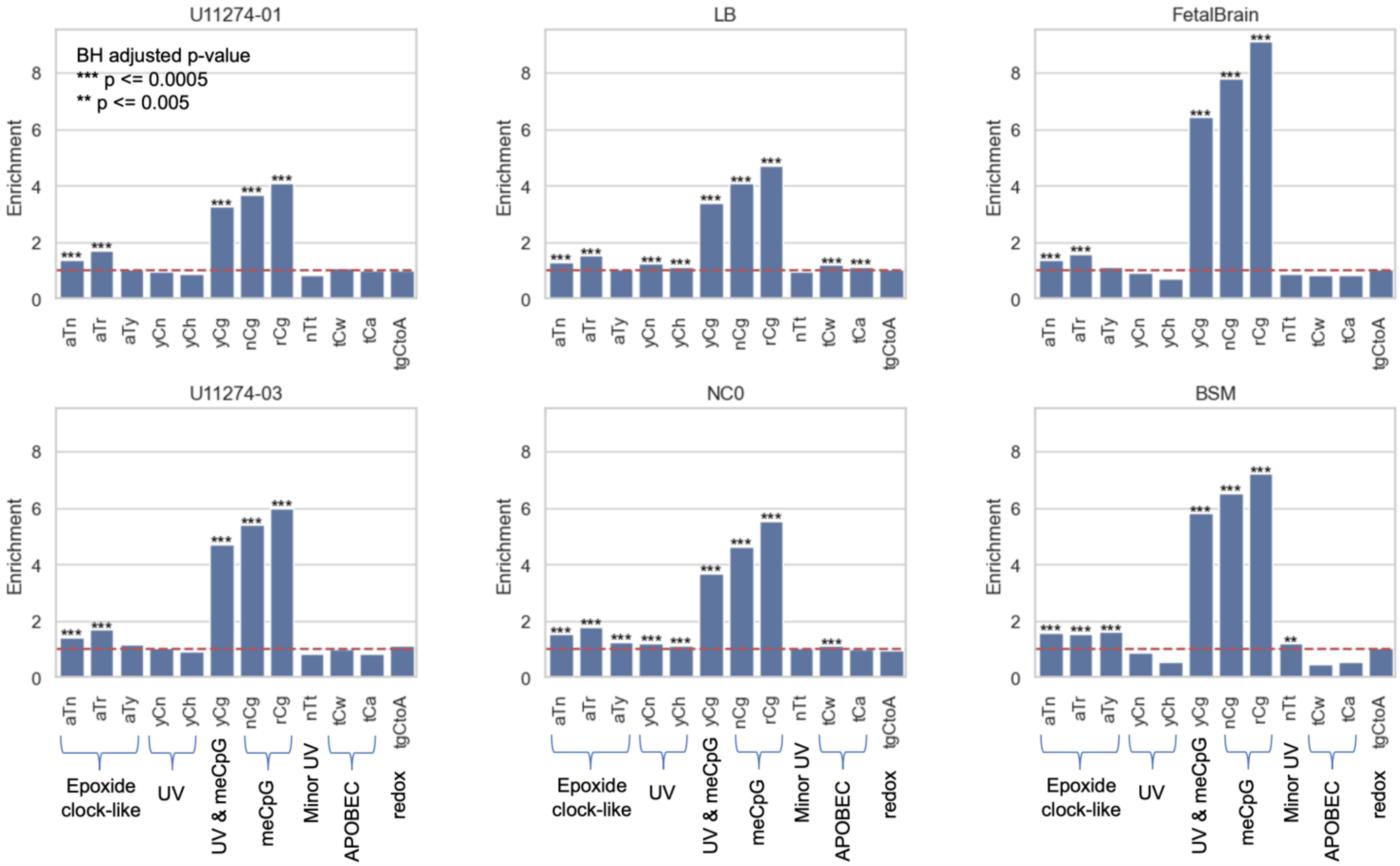


**Figure S5. Enrichment of trinucleotide mutational motifs across six different samples.** The fold enrichment of 12 pre-defined trinucleotide substitution motifs is shown across six samples: U11274-01 and U11274-03 (iPSC lines derived from urine), LB and NC0 (iPSC lines derived from skin fibroblasts), FetalBrain (single neuronal progenitor cells from fetal brains), and BSM (adult brain tissues). Fold enrichment represents the ratio of the observed mutation frequency within a specific trinucleotide context to its expected frequency based on local genomic sequence composition. The motifs are grouped by their association with known mutational processes: epoxide clock-like, UV, meCpG, APOBEC, and redox. Statistical significance assessed by Fisher's Exact test (see **Methods**) is indicated by asterisks above the bars, where *** represents $p \leq 0.0005$ and ** represents $p \leq 0.005$, with p-values adjusted using the Benjamini-Hochberg (BH) method to control for multiple testing. The red dashed line represents no enrichment (fold change = 1). **Table S2** provides detailed definitions of all trinucleotide motifs analyzed.

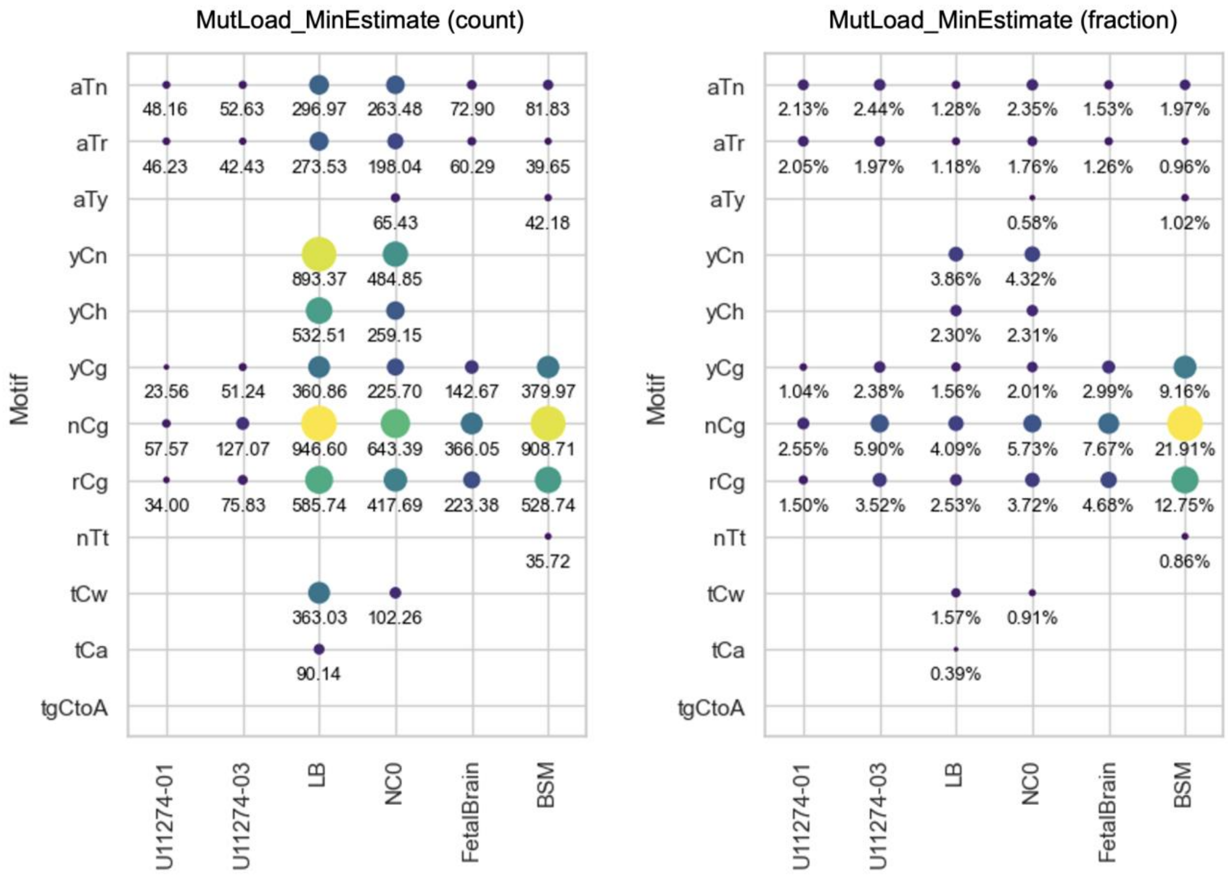


**Figure S6. Minimum mutation load estimates of 12 mutational motifs across six different samples.** Minimum mutation loads are shown in bubble plots for absolute counts (left) and fraction (right). Bubble size and color intensity correspond to mutation load magnitude, with specific values displayed below each bubble.

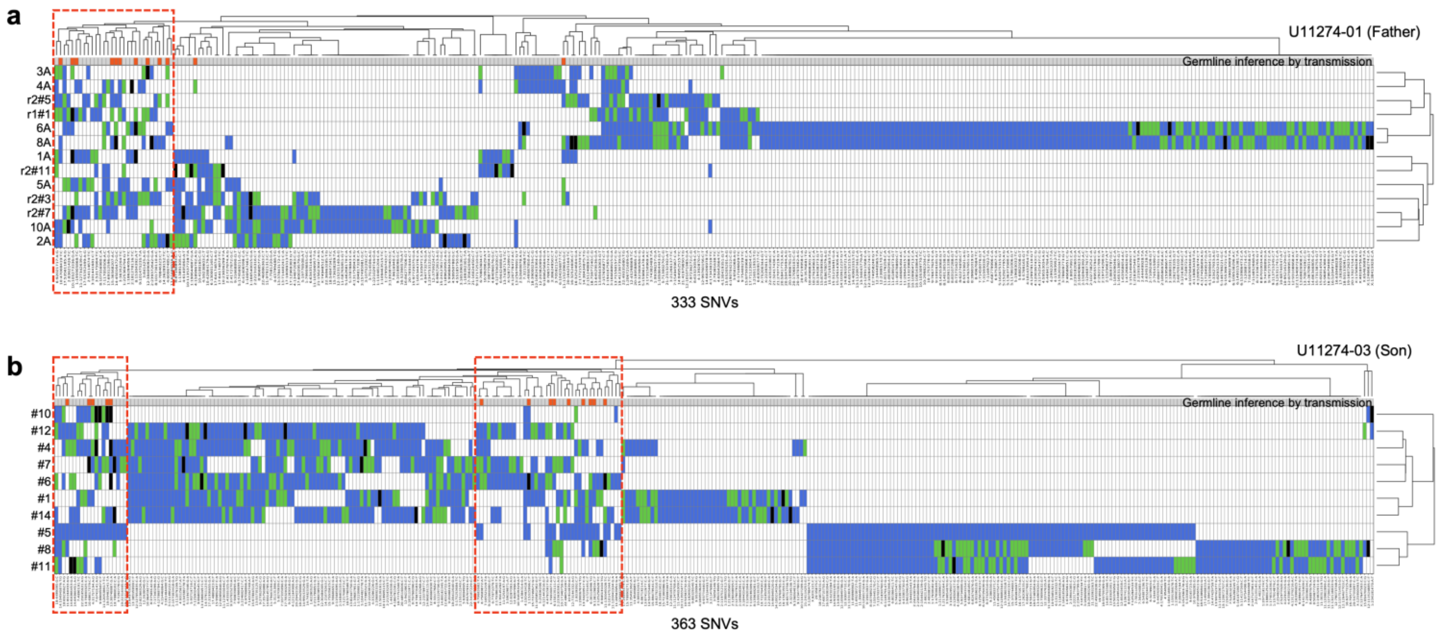


**Figure S7. Full version of mutational lineages including ambiguities from iPSC lines grouped by hierarchical clustering of shared SNVs. (a)** U11274-01 (father) and **(b)** U11274-03 (son). Founder cells of the iPSC lines are shown in rows, and shared mutations in columns. Colored cells in the heatmap indicate detected variants in each founder cell, with white cells indicating absence of variants. Blue indicates mutations detected in the discovery phase with at least three supporting alternative allelic reads. Green shows mutations rescued for lineage analysis with two supporting reads, while black represents those rescued with just a single supporting read. The top annotation bar marks germline variants (in red) inferred based on transmission between the father and son. Clusters enriched for germline variants (highlighted with red dashed boxes) were excluded from the lineage reconstruction shown in **Figure 3**.

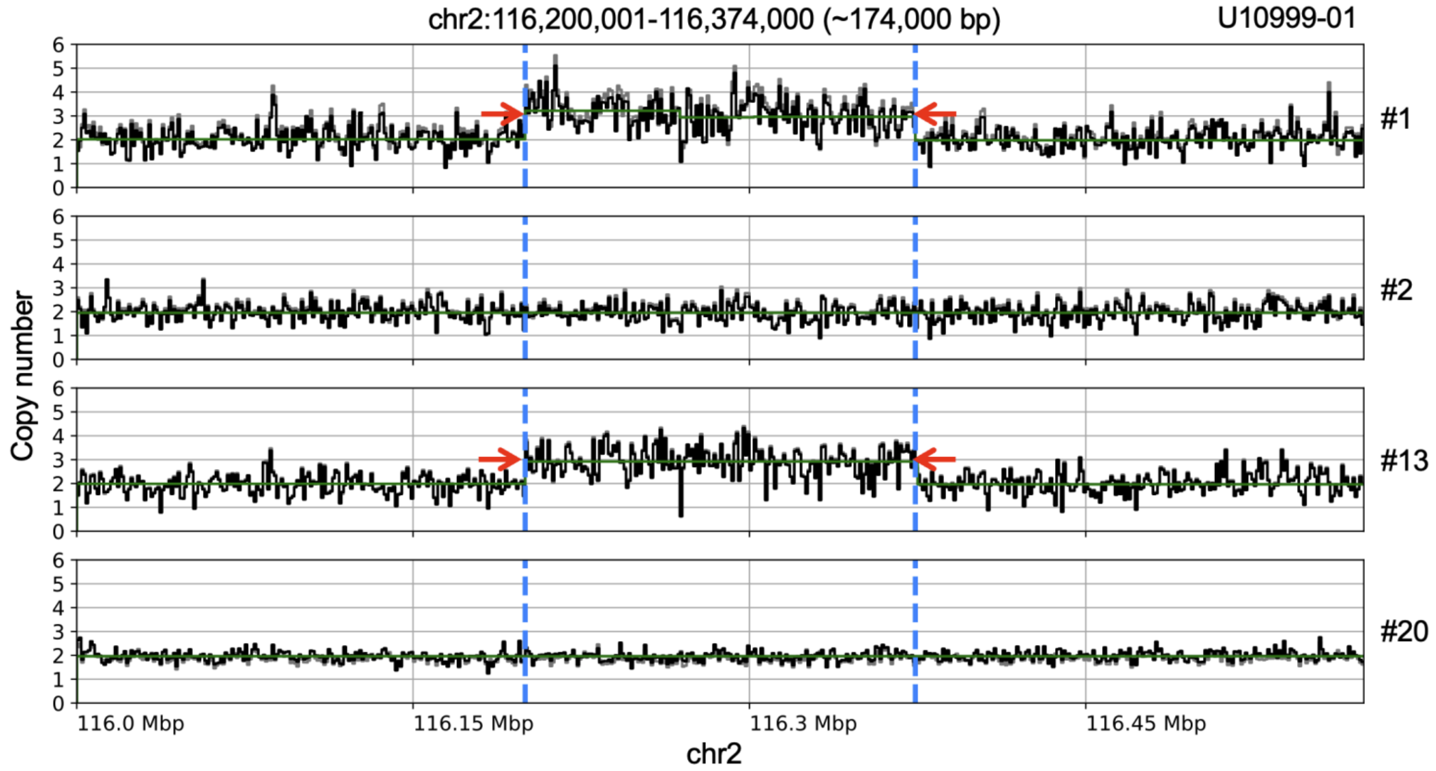


**Figure S8. Shared duplication on chromosome 2 identified in iPSC lines from individual U10999-01.** Copy number profiles inferred from read depth are shown for single cells. Cells #1 and #13 carry a somatic duplication (red arrows; ~174 kb) on chromosome 2, with the boundaries indicated by blue dashed lines.

### Supplementary Tables

**Table S1.** Summary of mutational motifs used in the knowledge-based motif-centered analysis.

| Motif | Ref. trinucleotide | Mutant trinucleotide | Knowledge | Hierarchy | Hierarchy note |
|---|---|---|---|---|---|
| aTn | aTn | aCn | Epoxide clock-like | all | all |
| aTr | aTr | aCr | Epoxide clock-like | partial | non-overlapping with yCn to yTn (UV) |
| aTy | aTy | aCy | Epoxide clock-like | partial | overlapping with yCn to yTn (UV) |
| yCn | yCn | yTn | UV | all | all |
| yCh | yCh | yTh | UV | partial | non-overlapping with nCg to nTg (meCpG) |
| yCg | yCg | yTg | UV and meCpG | partial | overlapping with nCg to nTg (meCpG) |
| nCg | nCg | nTg | meCpG | all | all |
| rCg | rCg | rTg | meCpG | partial | non-overlapping with yCn to yTn (UV) |
| nTt | nTt | nCt | Minor UV | all | all |
| tCw | tCw | tTw_or_tGw | APOBEC | all | all non-overlapping with nCg to nTg (meCpG) |
| tCa | tCa | tTa_or_tGa | APOBEC (preferred) | partial | non-overlapping with nCg to nTg (meCpG) |
| tgCtoA | tgC | tgA | redox | all | Part of SBS18 |

**Table S2.** Summary of validation results from experiments using PCR amplification coupled with Sanger sequencing.

| Mark in Fig. 3B | chrom | pos | ref | alt | Validated | Genotype by Sanger Sequencing | | | |
|---|---|---|---|---|---|---|---|---|---|
| | | | | | | #11 | #10 | #4 | #14 |
| *a* | 12 | 115751453 | G | C | O | **G/C** | G | G | G |
| *b* | 3 | 107223234 | T | C | O | **T/C** | T | T | T |
| *c* | 1 | 217630164 | C | A | O | **C/A** | C | C | C |
| *d* | 4 | 126506157 | G | T | O | **G/T** | G | G | G |
| *e* | 6 | 100382366 | A | T | O | **A/T** | A | A | A |
| *f* | 12 | 93007731 | C | T | O | **C/T** | C | C | C |
| *g* | 15 | 99892447 | C | A | O | C | C | **C/A** | **C/A** |
| *h* | 9 | 31283018 | G | C | O | G | G | **G/C** | **G/C** |
| *i* | 14 | 66145785 | A | G | O | A | A | **A/G** | **A/G** |
| *j* | 15 | 75505625 | C | T | O | C | C | **C/T** | **C/T** |
| *k* | 1 | 238802012 | T | G | O | T | T | T | **T/G** |
| *l* | 2 | 119751212 | G | T | O | G | G | G | **G/T** |
| *m* | 2 | 127607913 | C | A | X | C | C | C | unresolved |
| *n* | 10 | 114427209 | A | G | X | unresolved | A | unresolved | unresolved |
| *o* | 11 | 8546685 | G | T | O | G | G | **G/T** | **G/T** |
| *p* | 10 | 119985115 | C | T | O | C | C | **C/T** | **C/T** |
| *q* | 1 | 44240558 | G | A | O | G | G | **G/A** | **G/A** |
| N/A | 2 | 83975591 | C | T | O | C | C | C | **C/T** |

**Table S3.** List of copy number variations identified in urine-derived iPSC lines

| iPSC clone | Family | Relationship | Genomic coordinate | CNV type | Size |
|---|---|---|---|---|---|
| U10999-01_#1 | U10999 | Father | 2:116200001-116374000 | Duplication | 174,000 |
| U10999-01_#13 | U10999 | Father | 12:43958001-44257000 | Duplication | 299,000 |
| U10999-01_#13 | U10999 | Father | 2:116200001-116374000 | Duplication | 174,000 |
| U10999-01_#13 | U10999 | Father | 4:101017001-101452000 | Deletion | 435,000 |
| U10999-01_#20 | U10999 | Father | 17:9210001-9333000 | Deletion | 123,000 |
| U10999-03_#15 | U10999 | Son | 9:123681001-126331000 | Duplication | 2,650,000 |
| U10999-03_#30 | U10999 | Son | 11:114462001-114493000 | Deletion | 31,000 |
| U11274-01_1A | U11274 | Father | 1:219399001-219486000 | Deletion | 87,000 |
| U11274-01_2A | U11274 | Father | 13:64810001-65315000 | Deletion | 505,000 |
| U11274-01_3A | U11274 | Father | 9:17380001-17409000 | Deletion | 29,000 |
| U11274-01_3A | U11274 | Father | 11:112270001-114448000 | Deletion | 2,178,000 |
| U11274-01_3A | U11274 | Father | 11:114448001-135006000 | CN-LOH | 20,558,000 |
| U11274-01_5A | U11274 | Father | 2:153291001-153464000 | Deletion | 173,000 |
| U11274-01_6A | U11274 | Father | 16:83781001-83842000 | Deletion | 61,000 |
| U11274-01_8A | U11274 | Father | 16:83781001-83842000 | Deletion | 61,000 |
| U11274-01_10A | U11274 | Father | 11:9409001-9607000 | Deletion | 198,000 |
| U11274-01_10A | U11274 | Father | 4:73257001-73472000 | Deletion | 215,000 |
| U11274-01_r2#11 | U11274 | Father | 2:133767001-133782000 | Deletion | 15,000 |
| U11274-01_r2#11 | U11274 | Father | 6:36940001-37259000 | Duplication | 319,000 |
| U11274-01_r2#11 | U11274 | Father | 10:70960001-71113000 | Duplication | 153,000 |
| U11274-03_#1 | U11274 | Son | 3:67017001-72376000 | Duplication | 5,359,000 |
| U11274-03_#5 | U11274 | Son | 10:68467001-68591000 | Deletion | 124,000 |
| U11274-03_#10 | U11274 | Son | 16:78731001-78753000 | Deletion | 22,000 |
| U11274-03_#10 | U11274 | Son | 20:14683001-14864000 | Deletion | 181,000 |
